# Synchronization of mammalian motile cilia in the brain with hydrodynamic forces

**DOI:** 10.1101/668459

**Authors:** Nicola Pellicciotta, Evelyn Hamilton, Jurij Kotar, Marion Faucourt, Nathalie Degehyr, Nathalie Spassky, Pietro Cicuta

**Affiliations:** Cavendish Laboratory, University of Cambridge, Cambridge, United Kingdom; Cilia biology and neurogenesis, Institut de biologie de l’Ecole normale supérieure (IBENS), Ecole normale supérieure, CNRS, INSERM, PSL Université Paris, 75005, Paris, France

**Author notes:** NP performed experiments, data analysis and drafted the paper; EH performed numerical simulations; JK built experimental apparatus; MF, ND and NS provided numerous biological samples; NP, NS and PC designed the research; all authors contributed to writing the final paper.

**Keywords:** Motile cilia, Synchronization, multiciliated cell

## Abstract

Motile cilia are widespread across the animal and plant kingdoms, displaying complex collective dynamics central to their physiology. Their coordination mechanism is not generally understood, with previous work mainly focusing on algae and protists. We study here the synchronization of cilia beat in multiciliated cells from brain ventricles. The response to controlled oscillatory external flows shows that strong flows at a similar frequency to the actively beating cilia can entrain cilia oscillations. We find that the hydrodynamic forces required for this entrainment strongly depend on the number of cilia per cell. Cells with few cilia (up to five) can be entrained at flows comparable to the cilia-driven flows reported in vivo. Simulations of a minimal model of cilia interacting hydrodynamically show the same trends observed in cilia. Our results suggest that hydrodynamic forces between cilia are sufficient to be the mechanism behind the synchronization of mammalian brain cilia dynamics.

**Significance Statement:** It is shown experimentally, and also reproducing key qualitative results in a minimal mechanistic model simulated numerically, that in the motile cilia of the brain hydrodynamic forces of the magnitude that cilia themselves can generate are sufficient to establish the coordination of dynamics which is so crucial physiologically. This is the first experiment of its kind on multicilated cells, the key result is the unexpected importance of cilia num-ber per cell, with cells with fewer cilia much more susceptible to external flows. This finding changes the way in which we think about the question of collective cilia beating - it is not correct to simply examine isolated cilia and draw conclusions about the behaviour of cilia assemblies in multiciliated cells.

## Introduction

Fluid flow generated by a ciliated epithelium is a fascinating example of collective behaviour in nature: each single cilium beats in a non-reciprocal way along a defined direction and at the same frequency of the neighboring cilia, but with a different phase depending on the position. In many organs the result of this are “Metachronal waves” that are able to generate a stable flow parallel to the carpet of cilia (1). This dynamics has been reported to have fundamental roles in microorganisms (2) and in many organs of mammals. In the human airways, motile cilia generate flow for the clearance of mucus (3, 4). In the brain, the multiciliated ependymal cells covering all the ventricles ensure the cerebrospinal fluid circulation thought to be necessary for brain homoeostasis, toxin washout and orientation of the migration of newborn neurons (5). Defects in ciliary motility are the cause of many disease symptoms as bronchiectasis, hydrocephalus, and situs inversus (6). Although cilia coordination is required for the proper function of major organs, the mechanism behind it is still unclear.

The role of hydrodynamic forces during developmental phases has been demonstrated: To generate a macroscopic flow efficiently, motile cilia must beat all in the same direction. As shown first in the ciliated larval skin of *Xenopus* (7) and then in mouse brain (8), multiciliated cells were responsive to external hydrodynamic forces, and aligned their beating direction, when a physiologically relevant fluid flow was applied during development. Moreover, it was suggested (8) that the cilia-driven flow would act to refine the cilia beatings axis in a defined common direction after an initial orientation bias provided by a cell pre-patterning or an external fluid flow. After achieving alignment motile cilia beat at approximately the same frequency (with small cell to cell variability) all over the tissue, with a constant phase difference with respect to the adjacent cilia (9). It is the current hypothesis that cilia-driven flow could also itself be the mechanical origin of this coordination (10). From the pioneering studies of Huygens in the 1665, we know that oscillators with different intrinsic frequencies can reach synchronization if mechanically coupled (11). Cilia are microscopic filaments extended and moving into the fluid, and therefore the fluid will also act as a medium that will mechanically couple each single cilium. So, hydrodynamic interaction between cilia could provide the physical mechanism for their synchronization and metachronicity, (12). In the last decade, this hypothesis has been supported by simulations of pairs (13) and arrays of cilia (14–16), experiments with cilia models (17, 18) and with ciliated microorganisms from the genus *Paramecium* (19, 20*), Chamydomonas* (21) and *Volvox* (22, 23).

Despite many works supporting this view, recent experiments (24–26) showed evidence that disrupting the elastic coupling through the cytoskeleton also affected flagellar synchronization for the *Chlamydomonas* algae. The authors of (24) studied the phase locking between an external oscillatory flow and *Chlamydomonas* flagella. They measured that the required hydrodynamic forces for phase locking are over an order of magnitude larger than the ones between flagella, concluding that flagellar synchronization is due to elastic coupling. The importance of the flagellar inner coupling was also supported by theoretical studies (27, 28). This scenario enlightens the possibility that cilia coordination be achieved in nature in a organism-specific manner, thus leaving unclear what could be the underpinning mechanism in mammalian tissues.

Motile cilia in mammalian cells are (inevitably) coupled to some extent through the cilia-driven flow, but are also structurally connected through the actin mesh that links elastically all cilia of the same cell at the base (29–31). Even on the scale across confluent cells the epithelium could transmit forces elastically. In principle these connections could lead to synchronization (28), leaving the origin of cilia coordination in mammal organs still unresolved.

Inspired by the experimental setup in (24), we have tackled this problem by studying the response of single multiciliated cells to hydrodynamic oscillatory flows. We find that phase locking of the motile cilia with the external flow (entrainment) is possible, and that it strongly depends on the number of cilia per cell (Figure 1). Specifically, cells with few cilia (up to ten) are very responsive to external flow, and entrain with flow amplitudes comparable to the cilia-driven flows that regions of multiciliated epithelia can generate in vivo. By contrast, for cells with higher number of cilia per cell, entrainment required much higher flow velocities. We suggest that this is due to the hydrodynamic interactions between cilia in the same cell, and that multiple cilia can hydrodynamically screen the forces applied by the external flow, making the entrainment more difficult as the number of cilia increase. We support this view by quantitatively matching the data with a minimal hydrodynamic model. In the absence of external flow, we also measure an increase of the ciliary beating frequency (CBF) with the number of cilia, also consistent with hydrodynamically coupled cilia with coordinated motility. From these data we estimate the cilia-driven flow acting on each cilium. In view of the measured high susceptibility to flow, we find this cilia-driven flow to be quantitatively sufficient for sustaining hydrodynamic synchronization. Moreover we match the trend of our experimental data with simulations of a minimal model of cilia interacting hydrodynamically, showing the possible hydrodynamic origin of the measured effects. Overall our results suggest that hydrodynamic forces are sufficient to be the mechanism behind the synchronization of motile cilia in multiciliated cells of mammal brains.

**Fig. 1.**
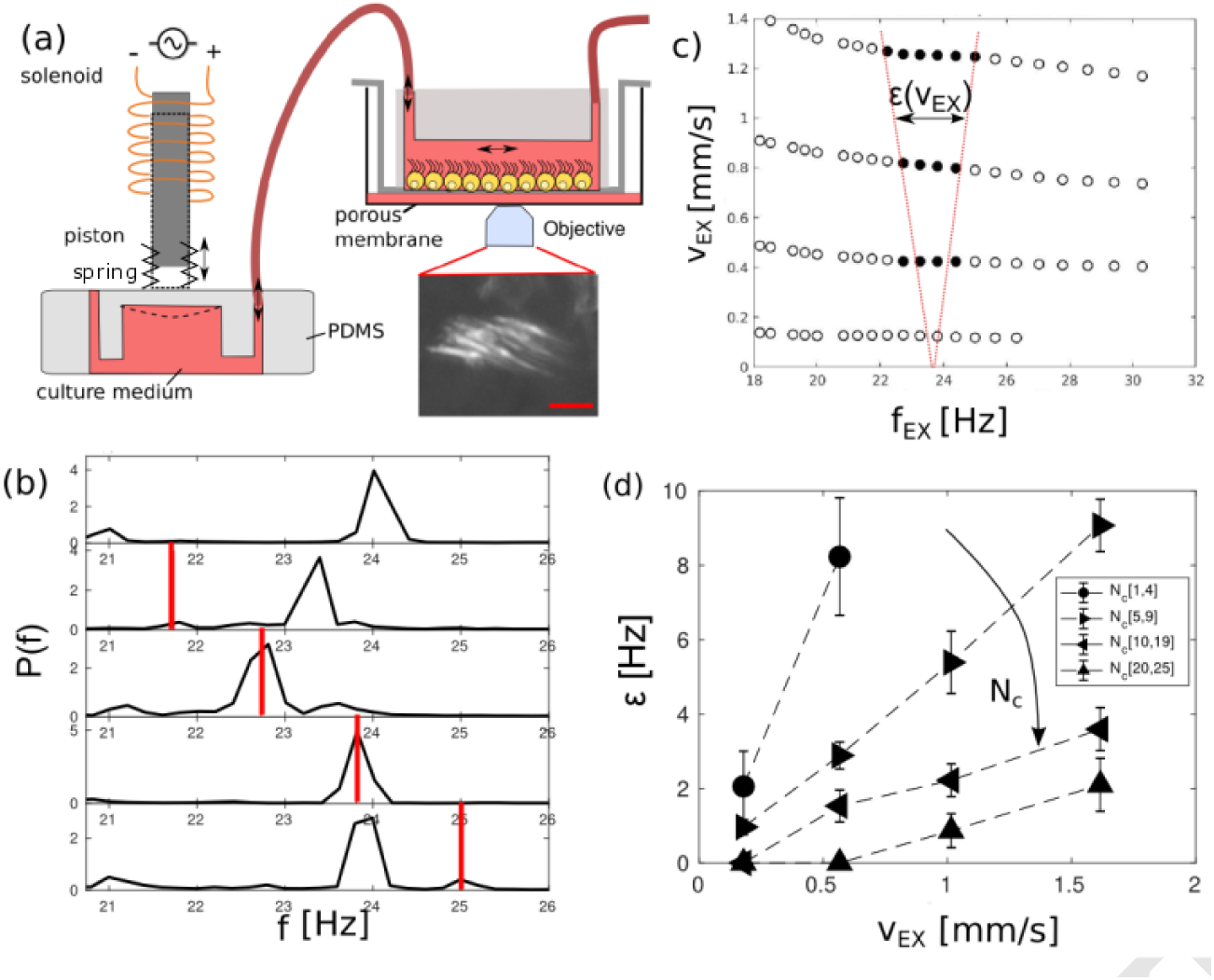
The strength of hydrodynamic coupling can be measured experimentally by applying an external oscillatory flow to a ciliated epithelium, whilst imaging cilia motility. (a) Schematic of the experimental setup, showing motility in a ciliated cell through a map of stan-dard deviation in time, scale bar 5 *µ*m. (b) The cilia beating frequency (CBF) is measured as the peak of the space-averaged periodogram of the cell pixels intensity, *P*^*S*^ (*f*). An entrainment event is identified when the peak of the *P ^S^* (*f*) coincides within 0.25 Hz with the frequency of the external flow *f*_*EX*_, (red line). The applied flow maximum velocity is *v*_*EX*_ = 0.6 mm/s. (d) Entrainment region, for the same multicilated cell, in the frequency and flow velocity domains; each marker here is the result of a separate recording. Markers are dark when the average CBF of the cell matches the frequency of the external flow. The entrainment strength, *∊*(*v*_*EX*_), is the range of frequencies spanned horizontally by the black dots. The entrainment region is highlighted here as the area between the red dashed lines; lines are a guide to the eye. (d) The en-trainment strength *∊*(*v*_*EX*_) increases proportionally to the applied external flow, but decreases with the number of cilia per cell *Nc*. Each line is the average of this trend over a group of cells with similar number of cilia. Complete datasets are shown in Figure S6.

## Results

### Experimental setup

We have used an *in vitro* culture assay of pure neural stem cells (NSC), isolated from neonatal mouse subventricular zone, that progressively differentiate into multiciliated ependymal cells (32). Cells were grown in a channel of PDMS attached to a plastic membrane (Corning Insert) that we call a “Transwell chip” (Figure 1a and methods). This Transwell chip culture, in a 6 wellplate with glass bottom, was imaged in Brightfield using a Nikon 60X 1.20 NA water immersion objective on an inverted microscope, and kept at 37°*C* and controlled *pH* using a custom made chamber (see methods). The inlet of the Transwell chip was connected to a custom-made fluid pump able to drive an oscillatory sinusoidal flow with defined frequency and velocity magnitude, see Figure 1a. We applied external flows spanning in frequency *f*_*EX*_ from 10 Hz to 35 Hz, and a flow velocity *v*_*EX*_ (average during half cycle) from 200 *µ*m/s to 2 mm/s. The lower value for *v*_*EX*_ was chosen to be close to the average net flow velocity created by ependymal cells in the brain (33). The applied range of frequency is related to the average ciliary beating frequency of the *in vitro* culture. As a calibration step, the flow created by the pump was measured from the trajectory of microparticle tracers at 7 *µ*m from the cell surface (see Methods). In each experiment a spatially isolated multiciliated cell was recorded for 5 seconds at 500 fps during each flow treatment to extract ciliary beating frequency (CBF) and phase. We took recordings only of cells close to the center of the channel, beating in the same direction of the applied flow and with a well defined peak in the frequency distribution (more information in Methods).

### Entrainment of multiciliated cells with external flow

We will refer to entrainment when we measure phase locking between the external flow and cilia within a cell, while the term synchronization will be used to indicate when many individual cilia coordinate their beating to a coherent collective beat pattern. The entrainment regimes for a multiciliated cell in the frequency and flow domains are shown in Figure 1c. Each marker represents a separate recording, the solid markers identify an entrainment event, when the measured CBF coincided with *f*_*EX*_, the frequency of the external flow, (more information in Methods). The regimes seen here agree with the shape of an Arnold tongue, a general property of coupled phase oscillators, and as was previously found for a pair of eukaryotic flagella of the *Chlamydomonas* algae under the effect of external flow (24) and in our model oscillator systems (34). From these data, one can define a entrainment strength *ϵ* (*v*_*EX*_) as the range of frequencies where we measured entrainment. For values of *v*_*EX*_ and *f*_*EX*_ within the entrainment area, the phase difference between the cilium and the phase external flow, Δ(*t*) = (*φ*_*cilium*_(*t*) − *φ*_*ϵX*_ (*t*))*/*2*π*, is constant over time (‘locked’) except for few occasional phase slips, Figure 2a-b. During a phase slip the cilia carry out one beat more (or less) with respect to the external force, and Δ(*t*) increases (or decreases) of one unit. By contrast, when *f*_*EX*_ and *v*_*EX*_ are outside of the entrianment area, Δ(*t*) increases dramatically with time (Figure 2a-b). This behaviour of Δ(*t*) resembles the general phase dynamics of a self-sustained oscillator under external forces and noise (35). Examples of the recordings that we used for the analysis are in Figure 2c and Figure S1 with link to video.

**Fig. 2.**
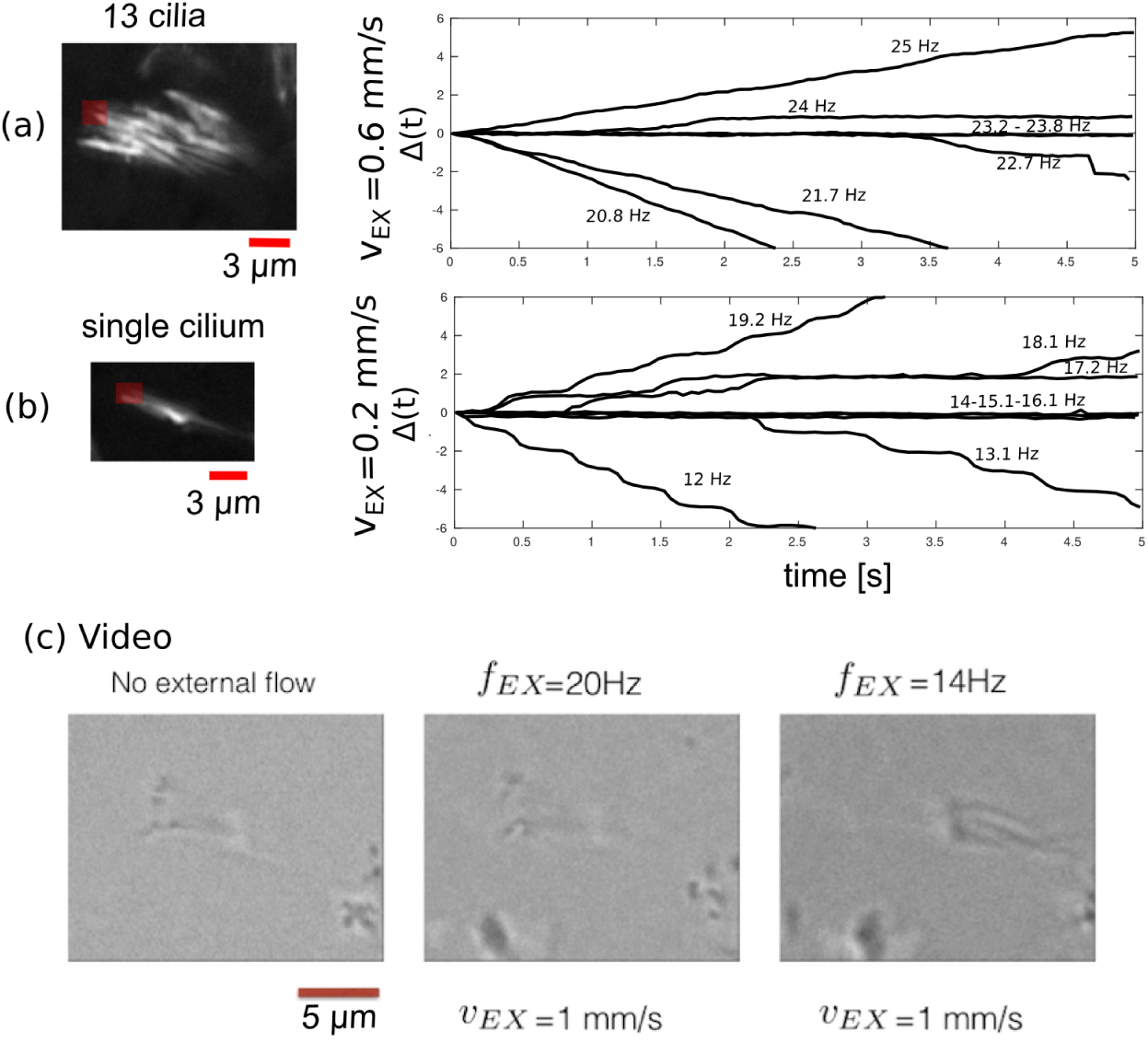
From high frame rate video microscopy record-ings we extract the frequency and phase of each cil-ium and the number of cilia per cell. (a) Phase dynam-ics Δ(*t*) when an external flow of *v*_*EX*_ = 0.6*mm/s* on a cell with 13 cilia. (b) Data for a single cilium when a ex-ternal flow of *v*_*EX*_ = 0.2*mm/s* is applied. The phase is constant (locked) when the cilium is entrained by the exter-nal flow. The interrogation windows used are marked in red over the standard deviation maps. (c) This example shows a movie of a single cell, being subjected to an external oscillatory flow. Images were acquired at 500 fps using a 60X objective, then analysed using background subtraction and spatial median filter of 3×3 pixels. http://people.bss.phy.cam.ac.uk/~np451/out/paper/video_singlecell.m4v

For each experiment the CBF was measured using a spatial average over an area segmented over the multiciliated cell of the time-domain Fourier Transform (FFT) spectra of pixel intensity, similarly to standard cilia CBF analysis (36, 37). The cilia phase dynamics Δ(*t*) can be found by tracking the passage of one cilium of the multiciliated cell in an interrogation window close to the position of the recovery strokes as was done on algae (22), more information in Methods.

### Entrainment depends on the number of cilia

Ciliated cells in culture (probably in contrast to most ciliated epithelia *in vivo*) are not always fully ciliated, and they exhibit varying number of cilia per cell (see Figure S8). We benefited from this situation and gathered results from 58 isolated multiciliated cells with different number of cilia. For each multiciliated cell we measured the entrainment strength *ϵ* as a function of the applied flow velocity *v*_*EX*_. We found that *ϵ* increases almost linearly with *v*_*EX*_, consistent with a recent study on the eukaryotic flagella of *Chalmydomonas* algae (24), Figure 1d. Moreover, *ϵ* does not change significantly with the intrinsic (absence of external flow) CBF of the cells (Figure S4), but dramatically decreases with the number of cilia per cell, see Figures 1d. The linearity of *ϵ* with the external flow *v*_*EX*_ and its trend with the number of cilia are highlighted in Figure 3b by showing the entrainment strength divided by applied the external velocity (*ϵ*(*N*)*/v_EX_*) as a function of *N*, the number of cilia per cell. Markedly, cells with 1 to 5 cilia were entrained by the external flow over a frequency range of around 2 Hz with flow velocity *v*_*EX*_ ≈ 200 *µ*m/s. In sharp contrast, cells with higher number of cilia were entrained over a detectable frequency interval only for strong flows *v*_*EX*_ *> ϵ* 600 *µ*m/s.

**Fig. 3.**
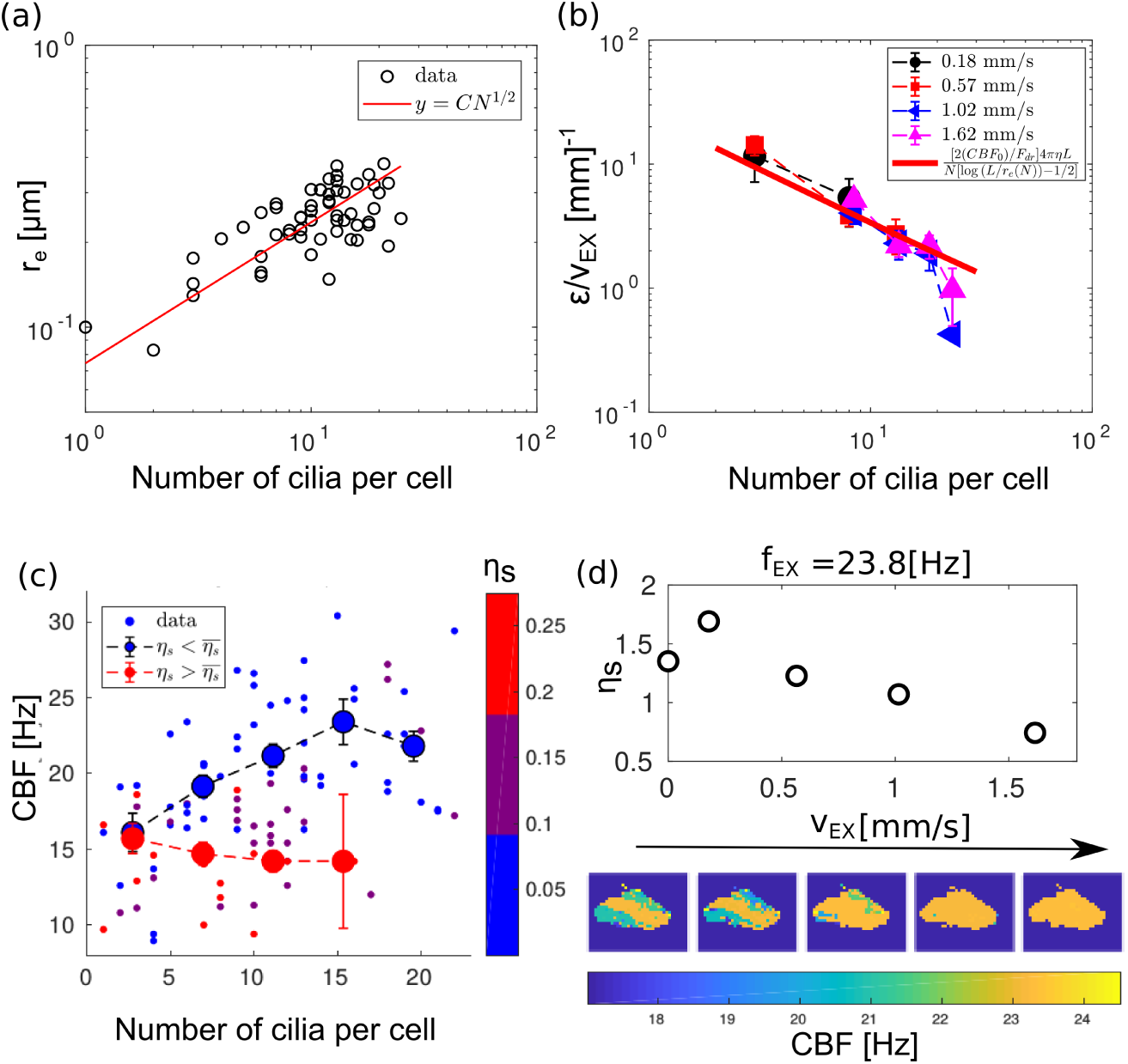
How a cell is affected by an external flow depends on the number of motile cilia: the entrainment of cilia motility gets weaker increasing the number of cilia, in agreement with a model where the external forces are hydrodynamically screened by the nearby cilia. (a) The effective radius of motile cilia scales with the square root of cilia number, with the prefactor compatible with the size of a single cilium; this supports the model that a cluster of cilia behaves hydrodynamically as an impenetrable rigid rod with an effective radius *re*, which contains all the cilia as a single bundle. For a cell with *N* cilia the effective radius is well approximated with *re* ≈ *CN* ^1*/*2^, with *C* = 0.08 ± 0.02*µ*m. (b) The measured entrainment strength decreases with the cilia number and collapses onto a single set when divided by the applied the exter-nal velocity (*c*(*N*)*/ v_EX_*). The trend with the number of cilia is well matched with the proposed model, using a single fit parameter *F*_*dr*_. The expression found in (a) is used for *r*_*e*_(*N*), with *L* = 11*µ*m and medium viscosity *η* = 10^−3^*P a s*. (c) In the absence of external flow, the ciliary beating frequency (CBF) increases with the number of cilia, but only for cells with low spatial noise *ηs* (i.e. cells where the cilia are already in sync). The data shows the binned medians for cells with low (blue) and high (red) spatial noise, highlighting the different trends in these two cases. Data were gathered from 120 cells. (d) The spatial noise *ηs* decreases when cilia are entrained by an external flow. This is shown with one cell as an example: each marker of spatial noise was calculated using a frequency map, illustrated in the panels at the bottom of the figure, with the colour corresponding to the measured frequency in each 4×4 pixel box.

This observation that high forces are required to entrain larger clusters of cilia is not in contrast with hydrodynam-ics because the external forces on each cilium could be hy-drodynamically screened by the forces applied the beating cilia nearby. This effect can be estimated: for an isolated cilium, the total drag force *F*_*EX*_ exerted by the external flow is proportional to the drag coefficient for a rigid rod *γ*= 4*πηL/*[log (*L/r*) − 1*/*2] (38), with *r << L* being respec-tively the radius and the length of the cilium and *η* the viscosity of the medium. Then, the external drag is *F*_*EX*_ = *γ v_EX_*. As the number of cilia increases, the cluster of dense cilia behaves hydrodynamically as an impenetrable rigid rod with length *L* and an effective radius *r*_*e*_(*N*) > *r* that contains all the *N* cilia, (39). The hydrodynamic screening by the nearby cilia can be seen as reduction of the effective drag 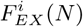 experienced by each cilium (*i*) in the cluster of *N*:

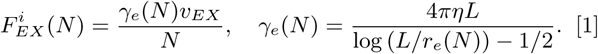

The effective radius *r*_*e*_(*N*) can be quantified approximately for each cell using the top view recordings of ciliary motion. We measured the area spanned by the cilia (*S*_*e*_) by counting the number of pixels where cilia motion was detected. The effective radius was then 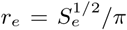 and normalised such that *r*_*e*_(*N* = 1) = 0.1*µ*m, equivalent to a radius of a single cilium (29). We found these data to scale as *r*_*e*_(*N*) ≈ *CN* ^1*/*^2 with *C* = 0.08 ± 0.02*µ*m, see Figure 3a, so compatible with very packed cilia, validating the former assumption of hydrodynamic impermeability for the cluster of cilia.

An exact solution that relates the entrainment strength to the external forces may be dependent on the intricate molecular response of the individual motors within a cilium (40) and on the efficiency of the ciliary beat (41). Here we use the known results for a minimal cilia model composed of a two state oscillator in viscous liquid driven by linear potential and entrained by an external periodic force (34). The molecular motors acting on the cilium are coarse-grained with a driving constant force *F*_*dr*_ driving the beat of the cilium (18). Applying this model to our system we find that the entrainment strength has a linear relationship with the ratio of the driving and external force:

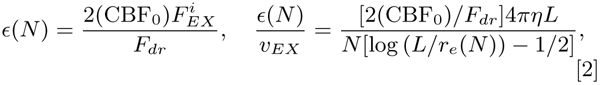

where CBF_0_ is the ciliary beating frequency of the isolated cilium. Since CBF_0_ and *F*_*dr*_ are intrinsic parameters of the isolated cilium, it is only the effective external force 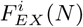 acting on the cilium that determines the trend of the synchronization strength with the number of cilia. The latter expression can be used to fit our data using only the free param-eter *F*_*dr*_, Figure 3b. For CBF_0_ we used the beating frequency of an isolated cilium, which in our culture was measured to be CBF_0_ ≈ 15 Hz, see Figure 3c, and the viscosity of the medium *η* = 10^−3^ Pa s. From the fit we found *F*_*dr*_ = 30 ± 3 pN. A rigid rod in the bulk of a viscous liquid, moving back and forward with amplitude *A* ≈ *L* ≈ 11*µ*m and frequency CBF_0_, would require a driving force 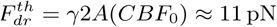. The fact that we estimated a slightly higher driving force is not consistent with the fact higher forces are required when translating a filament close to a surface (42). Despite its simplicity, the hydrodynamic model explains quantitatively the measured trend.

### CBF increases with the number of cilia per cell

In the absence of external flow, we expect that hydrodynamic forces between synchronized cilia in the same cell would also induce an increase of CBF. Previous theoretical work (14) predicted that arrays of beating cilia speed up their frequency, with respect to their intrinsic beating, when hydrodynamically synchronized, reaching a maximum increase when cilia are beating in phase. For 120 isolated cells we estimated the degree of synchronization using the frequency spatial noise *η*_*s*_ = *σ*_*f*_ */*(CBF), where *σ*_*f*_ is the standard deviation of the frequencies calculated on 4×4 pixels boxes over each cell, while the CBF is the median (frequency map in Figure 3d). We found that the CBF increases significantly with the number of cilia only for cells with low spatial noise *η* _*s*_ (high synchronization), Figure 3c. In this case, the CBF increases by roughly 50% for the cells with the largest number of cilia (*N*_*c*_ = [20 − 25]), compared to the ones with only few cilia. By contrast, cells with high noise do not show a significant CBF increase.

It is worth noting that the maximum CBF increase that we measured is much less sharp of what we expected for perfect synchronous beating cilia, when cilia in the same cell would behave hydrodynamically as a impenetrable object (Equation 1). This is not surprising in our culture, since cilia from the same cell beat with a finite frequency de-tuning or phase difference (metachronicity). By contrast, when cilia are entrained with the external flow, the spatial noise significantly decreases, Figure 3d, explaining qualitatively why the former assumption is valid during entrainment.

A similar CBF increase with cilia number was measured after cells were treated with actin drug Cytochalasin-D which progressively depolymerized the actin network connecting the centrioles in the cells and at the cell border (29, 31), supplementary Figure S5, suggesting that the observed CBF increase with cilia number is not due to cytoskeleton coupling.

By assuming that the CBF increase is due to the hydrodynamic forces between cilia, we can roughly estimate the magnitude of these forces acting on each cilium for a cell with low *η* _*s*_. In the previous paragraph, hydrodynamic interaction between cilia was represented by an the effective drag coefficient (Equation 1). However, the frequency detuning or metachronicity between cilia makes it more difficult to find an expression for the effective drag in this case. It is more convenient to write an explicit expression for the average of these forces during half cycle of the cilium, that we now define as 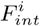, acting on the cilium *i* in a cell of *N* cilia. Using the former cilia model of a rigid cilium driven by a constant switching force, the velocity *v*^*i*^ of the cilium during half cycle is

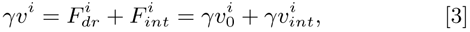

where 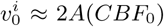 is the cilium av ge velocity in the absence of any interaction and 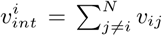 is the sum of the average flow velocity *v*_*ij*_ produced by the cilia *j* at the position of the cilium *i* during half cycle. If we consider a cell with large number of cilia, the CBF is increased by approximately 50%, so we can estimate the average velocity 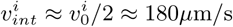.

We previously found that a similar magnitude of oscillatory external flow (*v*_*EX*_ ≈ 200 *µ*m /s) was sufficient to entrain small groups of cilia, and so certainly a single cilium (Figure 1d). We thus expect that the oscillatory cilia-driven flow 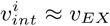 is also sufficient to ensure hydrodynamic entrainment of the cilium *i*. The same reasoning could be extended for each other cilium *j* in the cell, validating the hypothesis that cilia within a cell could be synchronized through internal hydrodynamic forces. Moreover, the magnitude of the interaction velocity 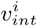 is in agreement with the measured flow field near eukaryotic beating cilia with similar length (22,43) and by the net flow generated by ependymal cells *in vivo* (33), supporting that the measured forces between cilia have hydrodynamic origin.

### Simulations of hydrodynamically coupled oscillators under external flow

In order to identify if the dynamics of cilia could stem from hydrodynamic coupling we compared our data with simulations of simplified cilia coupled hydrodynamically. We use a minimal rower model in which each cilium is represented by a single sphere driven by a geometrically updated potential, here each bead is driven by a repulsive harmonic potential, as illustrated in Figure 4a. This colloidal particle model, which we have been studying in the last ten years (18), is based on the physical intuition that, in a coarse-grained fashion, the degrees of freedom of the complex cilium shapes and activity can be captured by a rower’s driving potential. The main advantage of this approach is that it greatly simplifies the calculation of drag forces, both those acting on the individual object and the force induced by one object on another (44, 45). The bead moves away from the trap vertex until it reaches the switch point *A* + *x*_*s*_, where the trap is reflected and the bead reverses direction. This feedback-controlled motion of the potential is sufficient to induce sustained oscillations, and each particle undergoes long-time periodic motion with a fixed amplitude but free phase and period. Theoretical and experimental studies showed that pairs and chains of rowers can undergo synchronization (17) and even metachronicity (46, 47) when the separation is sufficiently small. To have similar scale to the biological system of this work, each cilium is simulated with a rower with radius *a* = 0.1 *µ*m, oscillating with fixed amplitude *A* = 7 *µ*m, tuning the strength of the parabolic trap to ensure an intrinsic frequency *f*_0_ = 15 Hz. These values were chosen to be similar to the ones that found experimentally tracking a single cilium (supplementary Figure S7). Simulations of chains were performed using a Brownian Dynamics code as in (44) varying the number of *N* rowers, keeping nearest neighbors separated on average by a distance *d*. We explored the set of *N* = [1, 2, 4, 6, 8, 10, 20, 30] and *d* = [0.4, 0.5, 0.6, 1.2] *µ*m (see Figure 4c). The rowers are coupled through the hydrodynamic forces via a Blake tensor, with the oscillations occurring at a fixed distance *z*_*wall*_ = 7 *µ*m above the no-slip boundary (44). For sufficiently small *d*, the chain of rowers synchronizes through hydrodynamic forces and beads oscillate at a common frequency *f* > *f*_0_, as seen in Figure 4c. As expected, the frequency shift *δf* = *f* − *f*_0_ initially increases with *N*, before saturating. The increase is more significant for rowers with small separation, due to the increase in their interaction strength. Interestingly, despite the simplicity of the rower model, the order of magnitude of *δf* in these simulations is the same as observed experimentally, although the latter is consistently lower. This could be explained by the fact that the distance between cilia in the same cell is much smaller (*d* ≈ 0.1 *µ*m) than the one that we can simulate (for distances *d* ≈ *a* near-field effects may play a role that we do not take into account).

**Fig. 4.**
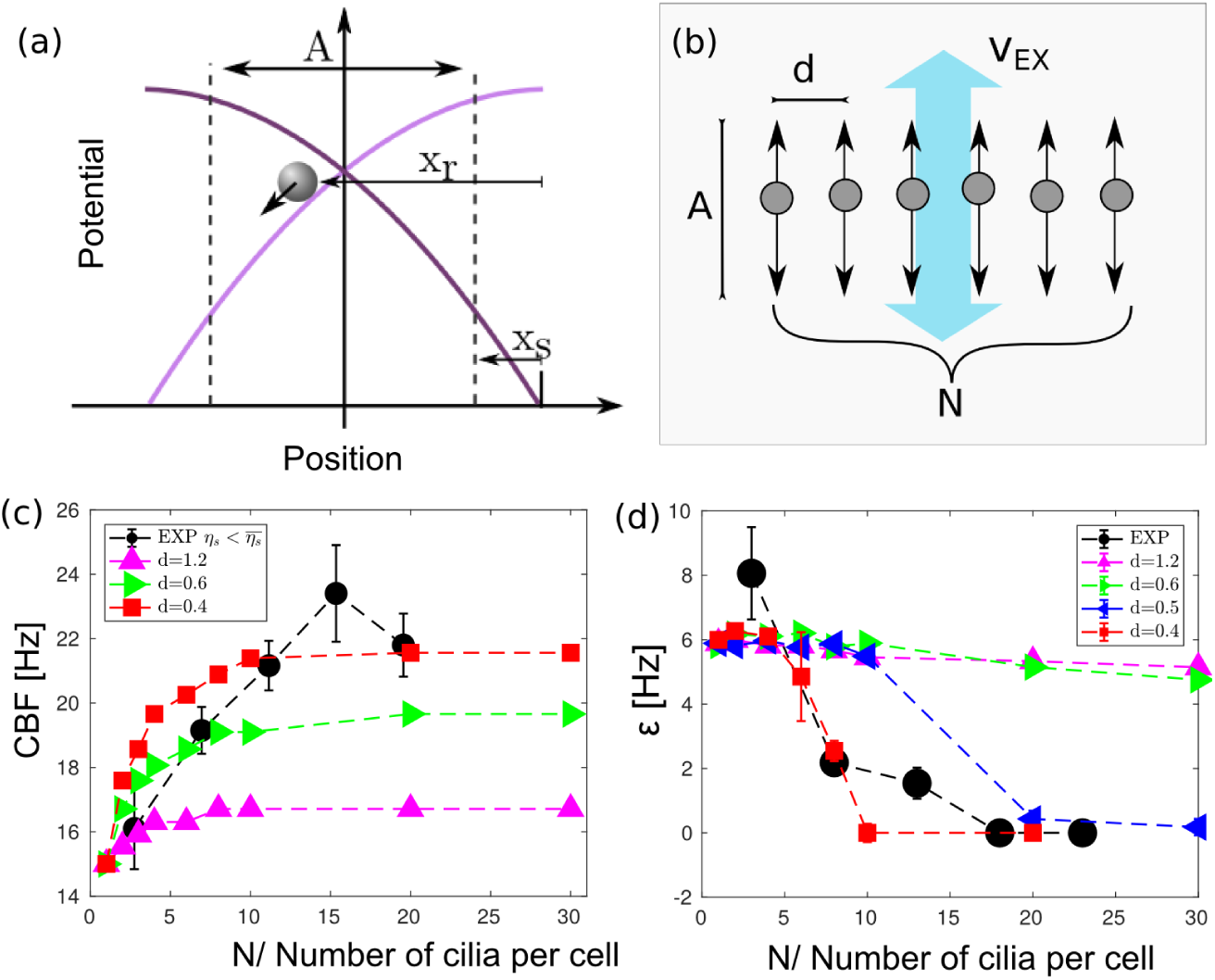
A coarse-grained cilia model where each cilium is a hydrodynamically coupled phase oscillator is studied numerically, and recapitulates the main trends seen in the experiment. (a) Geometric switch “rower” oscillator: Two potentials (purple and light purple curves) are switched on and off in turn using feedback loops. The other potential is switched when the particle reaches the position indicated by the black dashed line, maintaining a phase-free oscillation. (b) Illustration of the simulation rower geometry. (c) The median frequency of rowers increases with the number of rowers *N*. The growth is more marked as the rowers get closer, *d* = [0.4, 0.6, 1.2] *µ*m. The trend in simulations of the simple rower model matches with the experimental data (black points), where the distance between cilia is much smaller *d*_*cilia*_ ≈ 0.1 *µ*m. We only show experimental data from cells with low frequency spatial noise *η*_*s*_. Chain of rowers under oscillatory external flow. The entrainment strength *∊* measured with the external flow decreases as the number of rowers, or their density, increase. For the rowers with distance *d* = 0.4 *µ*m entrainment was not observed when the number of rowers was *N* > 10. In black are plotted the experimental data showing a similar trend, although the magnitude of the external flow used experimentally was an order of magnitude higher than the one used in simulations, and the distance between cilia is lower (external flows 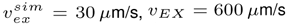)

We then simulated the chain of rowers under an external flow with a magnitude of 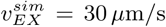. The chain was defined as entrained when more than 80% of the rowers were phase-locked with the external flow. Results are shown in Figure 4d. We observe a decrease in the entrainment region as the number of rowers in the chain increases. This effect is more pronounced when the density of the rowers increases. In the most extreme cases the chain does not entrain with the flow, which is consistent with the experimental data. It is worth noting that, for a given velocity, the smaller chain of rowers is entrained for a much larger range of frequency *ϵ* than what is observed experimentally, i.e. the susceptibility of cilia to an external flow is overestimated by the rower model. The simplicity of the model means it does not make sense to try and quantitatively match the experimental data, however it captures well the qualitative trends observed in the cilia.

## Discussion and conclusions

In the present work we combine experimental and theoretical approaches to understand the mechanism that leads cilia synchronization in the single multiciliated cell. Each cell has a variable number of motile cilia that are able to reach synchronization in frequency through an unknown mechanism of coupling. Studying this coupling is useful for the comprehension of the large scale problem, where thousands of multiciliated cells arranged in packed epithelia display metachronal coordination. In the present study we support experimentally that hydrodynamic coupling between cilia in ependymal multiciliated cell in the brain are sufficient to induce synchronization.

Firstly, we measured the hydrodynamic forces required for entrainment of cilia in isolated multiciliated cells. We found that multiciliated cells with few cilia (from 1-10) are very responsive to external oscillatory flow and can be entrained by a fluid flow around *v*_*EX*_ = 200*µ*m/s, similar in magnitude to the one that a cilium can generate during its power and recovery strokes (22) and to the total net flow generated by ciliated cells in the brain (33). We showed that the entrainment strength of a cilia bundle decreases with their number as we would expect for hydrodynamically coupled cilia. The forces applied by the external flow on each cilium are hydrodynamically screened by the beating cilia nearby, so each cilium fells a lower drag than when isolated. This trend is quantitatively described by a model that assumes the group of packed cilia to behave hydrodynamically as an impenetrable rigid rod with an effective radius (that depends on the number of cilia) and each cilium beating driven by an average constant force, (34).

Secondly, we measured that multiciliated cells with higher number of cilia were in average beating with larger CBF in a way that depended on the cilia frequency mismatch in the cell. The CBF increase was not affected after depolymerization of the cell actin network, supporting the hypothesis that this effect has not cell cytoskeleton coupling origin. By contrast, a CBF increase is consistent with hydrodynamically synchronized cilia, (14). Arguing that CBF increase has only hydrodynamic origin, we estimated the cell-driven flow acting on each cilium in a synchronized multiciliated cell. We found that this flow is quantitatively sufficient to entrain the motility of each cilium within a cell validating that hydrodynamic forces between motile cilia are sufficient to lead to synchronization.

To further verify the importance of hydrodynamic forces, we carried out simulations of motile cilia studying the effects of cilia number and external flow, within the minimal model of rowers. In this model, actively driven oscillators interact only through the fluid flow that they create themselves by oscillating. We found that the trends of the simulation results match the experiments on live cilia. Specifically, simulations predict an increase of the CBF with the number and density of cilia that is similar to the one measured. Moreover, they show a threshold with the number of cilia above which the entrainment of a chain of rowers is not possible anymore, as observed experimentally.

Biochemical or mechanical elastic coupling at the base of the cilia could also contributes to the synchronization of the cilia. However, in the algae *Chlamydomonas*, where the elastic coupling between cilia has been shown to be important for the synchronization (25, 26), hydrodynamic forces required for entrainment were measured to be an order of magnitude higher of the physiological ones (24). By contrast here we measure a great susceptibility of motile cilia to external flow in the same order of the cilia-driven flow reported in literature (33) and quantitatively in agreement with the cilia-driven flow that we measured in a synchronized multiciliated cell. These results, true at the single cell level, are expected to generalize to the ciliated epithelia in mammals brain, although *in vivo* other variables such as the cell orientation and density could play important roles for the establishment of global beating coordination (8, 48). Further experimental investigations are also required to validate these results for mammalian multiciliated cells propelling mucus, such as in the airways, where cilia beat in a non-Newtonian medium and are also tethered by mucins (49).

## Methods and Materials

### Transwell chip

PDMS substrates with channel features were bonded to Corning Transwell insert membrane using PDMS prepolymer as glue (50). PDMS prepolymer was mixed with curing agent at a weight ratio of 12:1. The mixture was then cast onto an aluminium mould having positive reliefs of the channels with 1 mm × 1 mm rectangular area and 12 mm long and cured at 60°C overnight. The mould was produced by CNC mill machining. The cured PDMS layer was peeled from the aluminium substrate and holes of 2 mm diameter were punched through the cured PDMS substrates at the end of the channels. Then PDMS prepolymer with (ratio 10:1) was spin-coated on a clean glass cover slide for 1 min (10 s at 500 rpm, ramped at 500 rpm/s to 3000 rpm for 60 s) to generate a thin mortar layer (51). This thin PDMS prepolymer serves as ‘mortar’ to strongly bind the PDMS substrate with the membrane. The PDMS substrates with channels were placed onto the glass slides spin-coated with the adhesive PDMS mortar and allowed to stay in contact for 30 s. Then the PDMS substrate were gently placed on Transwell Corning inserts with a polyester membrane 24 mm diameter and 0.4 *µ*m porosity. The combined pieces were left to cure at room temperature for at least 2 days in 6 well plates. Higher curing temperature would result in melting of the glue between the membrane and the insert. Then, two short pieces of tubing of 0.50” OD were placed at the inlet and outlet of the channels to facilitate the entrance of fluid using pipettes. Before seeding the cells, the complete Transwell chip was sterilized under UV for 30 minutes. PDMS offers many advantages for microfluidic cell culture because of its gas permeability, optical transparency, and flexibility, and has been used for the last decades for long term culture and cell differentiation of many cell types (52, 53). However, for long term culture in microfluidics, constant perfusion of media is needed because of the limited nutrients available in the small channel volume (54). For this reason, the PDMS channel was bound to a Corning Transwell insert, where the medium in the basolateral compartment acts as a nutrient reservoir that reaches the cells through the porous membrane. This allows the culture and differentiation of ependymal cells in a channel without the need of media perfusion. We found this extra medium to be a critical requirement for a good differentiation of the neural stem cells in the channel.

### Cell culture

All animal studies were performed in accordance with the guidelines of the European Community and French Ministry of Agriculture and were approved by the Ethics committee Charles Darwin (C2EA-05) and “Direction départementale de la protection des populations de Paris” (Approval number Ce5/ 2012/107; APAFiS # 9343). Differentiating ependymal cells were isolated from mouse brain and grown in flask as previously described (32). When cells were 70-80%confluent in T25 Flask, they were seeded at concentration of 10^7^ cell/ml in the Transwell-chip in 10% FBS 1% P/S DMEM medium. The day after, the cells were rinsed with PBS, and medium was switched to 1% P/S DMEM medium. Medium was not changed during all the period of differentiation. After 15 days cells were fully differentiated. In the culture a small percentage of multiciliated cells has motile cilia, lacking a common beating direction and frequency. For our analysis we focused on 58 multiciliated cells from 8 different Transwell inserts, with all the cilia beating in a defined beating direction and showing a single peak spatial frequency distribution. More info on the statistics in supplementary Figure S3.

### Solenoidal pump and flow calibration

The inlet of a Transwell insert is connected through tubing to a PDMS chamber fully filled with fresh medium. An alternating flow of medium is created when the top membrane of the elastic chamber is moved up and down by the piston. The oscillations of a soft iron piston were controlled by the magnitude and frequency of alternate current (AC) flowing into a solenoid copper coil around the piston Figure 1 and a preloaded spring in a similar fashion to commercially available solenoidal fluid pump. The solenoid was made by wrapping copper wire (0.5 mm) 300 times around an aluminium case for the soft iron piston and the spring. The piston was tightly connected to the top membrane of the PDMS chamber through a screw ending embedded in the PDMS. The PDMS chamber is 5 mm height and with top area 10 mm × 10 mm. The alternate current was created by a function generator connected to a current drive source. The output current was monitored with oscilloscope through a current proportional voltage output. We tracked 1 *µ*m polystyrene tracers particles to measure the flow and frequency created by the solenoidal pump into the Transwell insert. The calibration was performed in a Transwell-chip without cell. Recordings of the oscillating particles were taken at 500 fps and with 40X and 20X objectives at distances of 7, 14 and 21 *µ*m from the wall surface. The average velocities during half period were measured by tracking the particle displacements. The minimum applied velocity was chosen to be similar to the flow measured in vivo by tissue ependymal ciliated cells of ≈ 200 *µ*m/s (33). The maximum applied flow velocity is 10 times the physiologically relevant one. Supplementary Figure S2 shows the output flow velocity varying the amplitude and frequency of the AC signal in the solenoid. The generated flow velocity was measured to be higher for low frequency due to the resonant response of the system.

### Image acquisition and analysis

Images were taken with a Ti-E inverted microscope (Nikon, Tokyo, Japan) equipped with a 60X WI objective 1.20 NA. High speed videos were recorded at 500 fps using a CMOS camera (model No. GS3-U3-23S6M-C; Point Grey Research/ FLIR). At each image was subtracted a running average of 60 frames. To define entrainment events, CBF is measured as the highest peak of the first harmonic of the average periodogram, 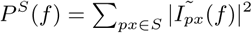, as in (36, 37, 55), with 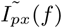 the FFT over time of the pixel intensity *I*_*px*_, and *S* the ensemble of pixels in a sub image thresholded for ciliary motion. The sub image was defined to include all pixels for which the standard deviation of the intensity over time is larger than a threshold value (this found with Matlab function using Otsu’s method), Figure 5. Entrainment with the external flow is identified when the frequency of the signal highest peak 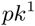 coincides with the frequency of the external flow within a certain error *σf*. For our acquisition rate *σ* = 0.25 Hz, the requirement is half of our frequency resolution. This method was implemented for high values of external flow *v*_*EX*_ > 1*mm/s*, when the oscillatory external flow induces a periodic defocusing of the field of view. This effect was detectable in the periodogram and often contributes to the highest peak of the signal, making it more tedious to measure CBF and entrainment events. For this reason, when the first peak of the signal 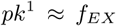, we also compared the second higher peak 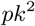 to the value of 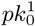, the signal peak in absence of flow. When 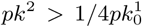, we identify such entrainment events as false, and the second peak as the real contribution from the cell to the signal. This process was found to be valid and stable for every movie that we inspected.

**Fig. 5.**
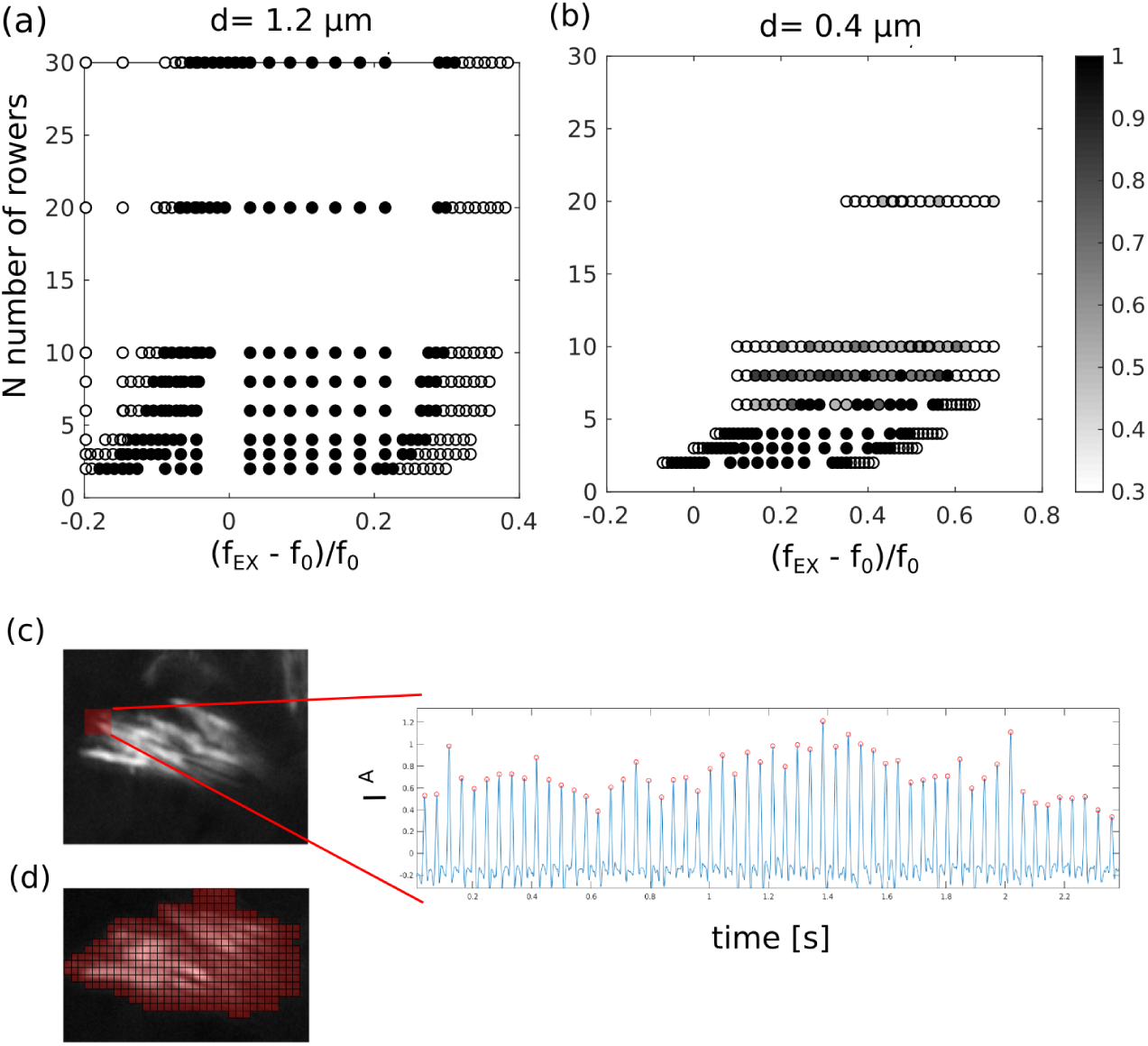
Simulations and image analysis methods. (a-b) Simulations: synchronization graphs in frequency domain as a function of the number of rowers, using external flow velocity *v*_*EX*_ = 30 *µ*m/s for both the two cases. The col-ormap represents the percentage of rowers synchronized in each simulation, 1 is when 100% of all the rowers are synchronised with the external force. (c) The pixel intensity *I*^*A*^(*t*) over time is used to identify the phase of a cilium or cilia within a cell. Each peak is the time when a cilium complete a beat cycle. The interrogation area is indicated in red on the standard deviation map. (d) The sub image used for the calculation of the CBF and defined using the standard deviation of the pixel intensities over time.

The phase dynamics was calculated by tracking over time the pixel intensity of an interrogation area *A*, chosen to be close to the recovery stroke of the cell, *I*^*A*^ = − ∑_*px∈A*_ *I*_*px*_. This area is small, such that only one or at most two cilia can be detected per time. When a cilium passes through this area, the pixel intensity *I*^*A*^ spikes, see Figure 5. The time of the spike is an indication that the cilium completed a cycle, and the phase of the cilium *φ*_*cilium*_ increases by 2*π*. The number of cilia per cell was counted by inspection of movies at 500 fps, after median filtering (window of 3 × 3 pixels) and contrast correction. All the analysis where performed in Matlab.

### Simulations

The simulation scheme builds on previous work (34, 44, 45) but implements external flow. Rowers are driven by repulsive harmonic traps, with the potential 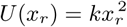. The traps are updated when *x*_*r*_ > *A* + *x*_*s*_, with the switch condition *x*_*s*_ = 1 *µ*m and the amplitude *A* = 7 *µ*m. The beads themselves have a radius *a* = 0.1 *µ*m and their position is updated via a Langevin equation,

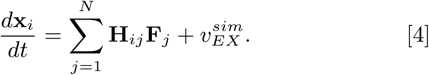

The rowers are coupled through the Blake tensor **H**. The tensor is implemented in a plane at a constant height above the no-slip boundary, *h* = 7 *µ*m (44). The drag is *γ*= 6*πηa*, where *η* = 0.0022Pa*·*s.The external flow 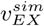 is implemented as a square wave, it has an amplitude *v*_*e*_ and it changes direction every *N*_*f*_ frames. The switching results in a period *N*_*f*_ *δt* s, where *δ t* is the simulation time step.

The simulations were run for 10 s (150 cycles), with time step *δt* = 3 · 10^−5^ s (9 · 10^−4^ cycles). Each case was repeated 10 times, with the initial positions of each rower within the trap drawn from a uniform distribution 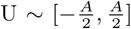. The entrainment strength *ϵ* is calculated for each seed, and the average reported here.

A rower is considered entrained to the external flow if after the initial transient period (3 s) fewer than 5 phase slips occur. When more than 80% of the rowers in the chain are entrained, the system is considered phase-locked. The rowers near the edge of the longer chains are not included in this calculations, due to edge effects stemming from having fewer neighbors. Most entrainment strengths *ϵ* are calculated by finding the difference in detuning between the smallest and largest values Figure 5. For some of the high density cases, phase-locked and non-phase-locked states were too interspersed to use this approach. Instead, *ϵ* was calculated by summing the number of states recorded as phase-locked and scaling by the frequency sampling resolution 0.017*f*_0_.

## ACKNOWLEDGMENTS

NP and PC were supported from the European Union’s Horizon 2020 research and innovation program under the Marie Sklodowska-Curie grant agreement 641639 ITN BioPol, PC and JK also supported by ERC CoG HydroSync and EH by Cambridge Trusts. MF, ND and NS were supported by INSERM, CNRS, École Normale Supérieure (ENS), ERC CoG grant 647466.

## Supplementary materials of: Synchronization of mam-malian motile cilia in the brain with hydrodynamic forces

**Fig. S1.**
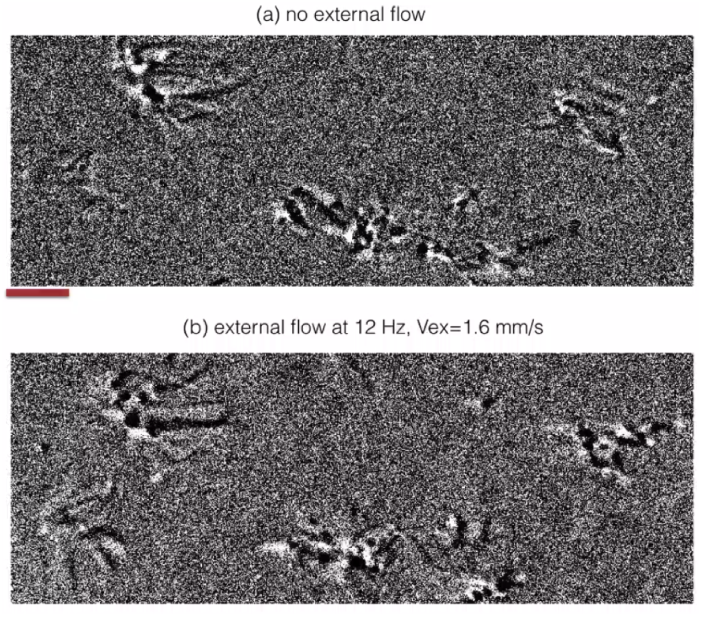
Very strong external flow can synchronise a group of cells to beat in synchrony. (a) Ciliated cells beating without external flow. (b) External oscillatory flow of *u*_*EX*_ = 2 mm/s at *f*_*EX*_ = 12 Hz. This strong flow induces frequency synchronisation and also alignment of beating direction of some cells that were beating misaligned to the external flow. The original direction of beating was recovered after the flow stopped. Images were acquired at 500 fps using 60X objective, then analysed using background subtraction, contrast enhancement and spatial median filter 3 × 3 pixels. Videos are played at 50 fps, 10 times slower the original recording. http://people.bss.phy.cam.ac.uk/~np451/out/paper/video_cellgroup.m4v

**Fig. S2.**
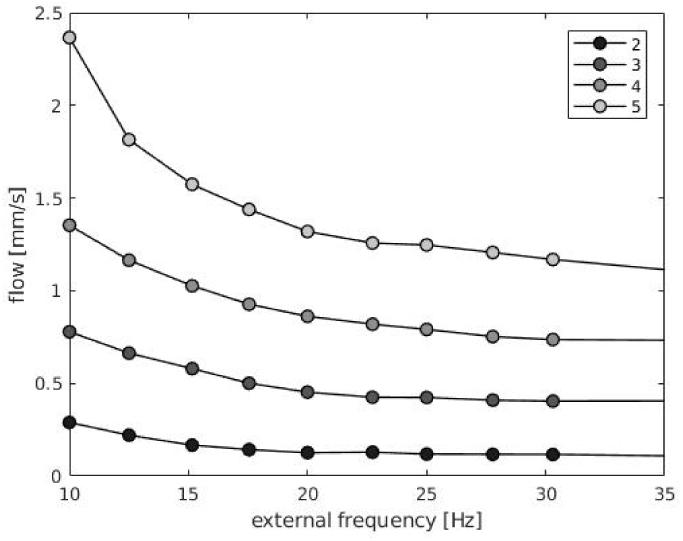
Flow created at the center of the transwell insert when a defined current was applied at the ends of the solenoidal pump, the flow as been calibrated using 1 *µ*m beads at the defined distance above the surface of [7, 14, 21 *µ*m] in order to have an accurate measure of the flow within the channel. The graph shows the value at 7 *µ*m from the wall. These values have been used for the graph reported in the main text.

**Fig. S3.**
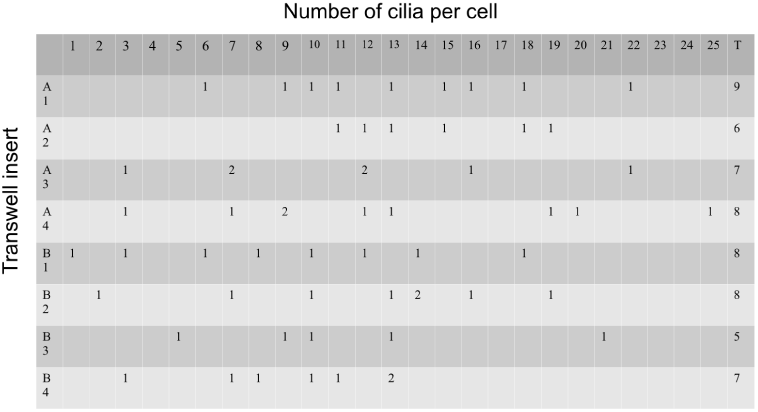
We used a total of 58 cells from 8 different Transwell inserts cultures. For each insert we imaged from 5 to 9 multiciliated cells under flow. The table gives information for each cell about the number of cilia, and in which Transwell insert it was cultured. Inserts (A1 to A4) were ependymal cultures 2 weeks before the B inserts (B1 to B4).

**Fig. S4.**
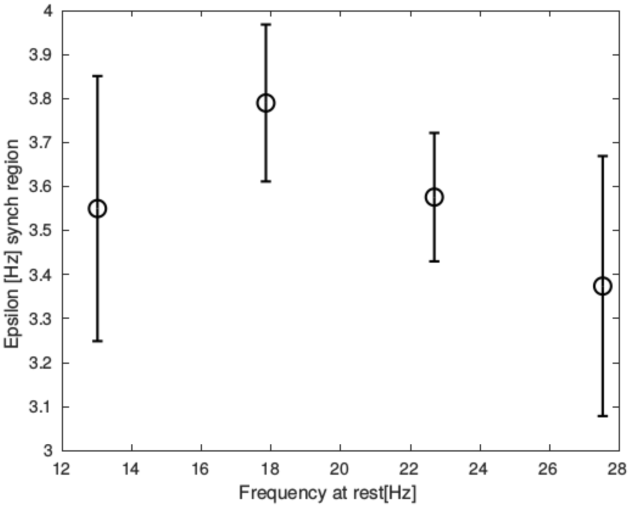
The synchronisation strength *∊* of cilia within a cell does not significantly depend on the intrinsic cilia beat CBF. The graph shows *∊* as a function of the intrinsic CBF of the multiciliated cells when an external flow of *u*_*EX*_ ≈ 1 mm/s is applied.

**Fig. S5.**
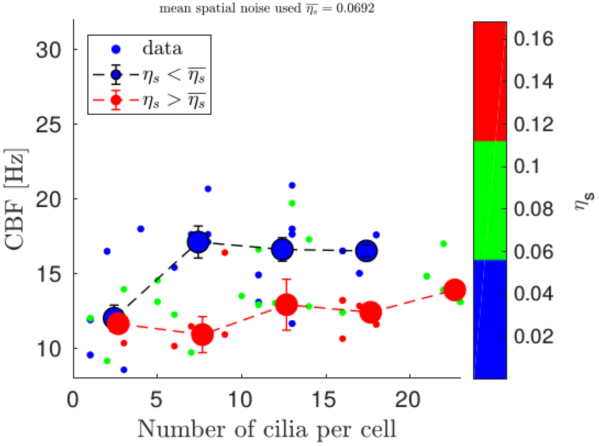
CBF vs cilia number for cells treated with actin drug Cytochalasin-D (80 cells from 3 treated samples). CBF increase significantly for cell with low spatial noise *η*_*s*_. Comparing cells with few cilia (*N* ≈ [1 − 5]) with the ones with many cilia (*N* ≈ [15 − 20]), the increase in CBF is around 50% (from 11Hz to 17Hz), similar to the one that we measured for not treated cells. It is worth noticing that the CBF generally decreased of few Hz respect to the control (treated with DMSO). Both control cells with DMSO and cells treated with 2uM Cytochalasin-D were stained in 4% PFA for 10 minutes, permeabilised with Triton x-100 0.1% in PBS, and incubated for 1 hour with Nucblue R37605 (1 drop for mL of PBS) and with phalloidin for actin (Sir-Actin, 0.2 uM) following proprietary protocols. Z-stack were taken with confocal microscope (slices of distance of 0.15um each) and can be found at the following link: DMSO treated cells (control) http://people.bss.phy.cam.ac.uk/~np451/out/paper/control_DMSO_52um_z2.4um.avi, and Cytochalasin-D treated http://people.bss.phy.cam.ac.uk/~np451/out/paper/cytoD_2uM_55um_z2um.avi

**Fig. S6.**
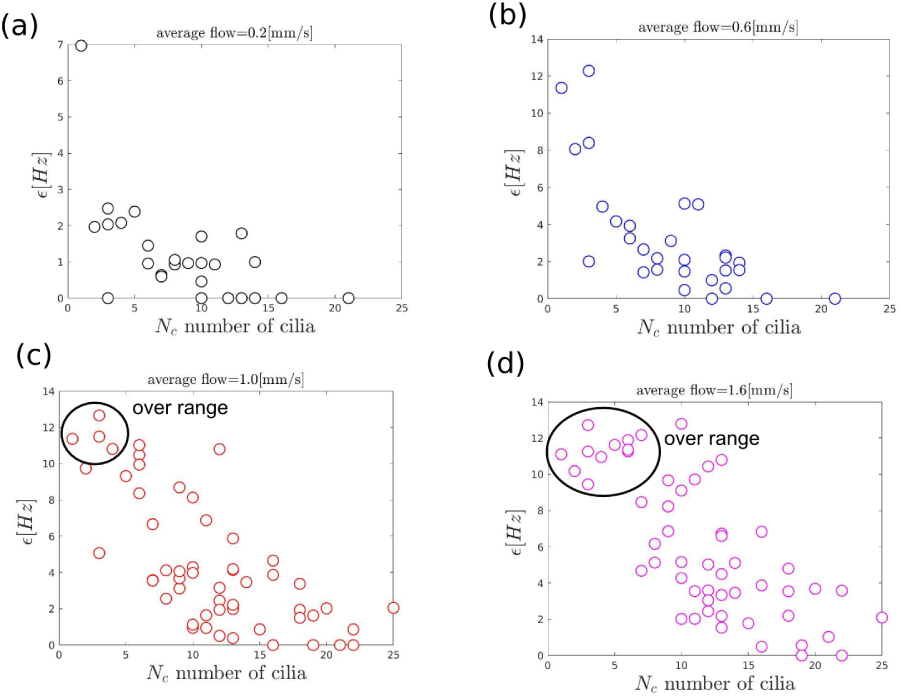
Synchronisation strength *∊*(*u*_*EX*_) for each cell as a function of the number of cilia per cell *N*_*c*_. Each plot shows data using a different external flow average velocity *u*_*EX*_ (shown on top of the graph). The points in the “over range” circle correspond to the cells that were entrained for the whole range (or nearly whole) of frequency spanned by the external flow.

**Fig. S7.**
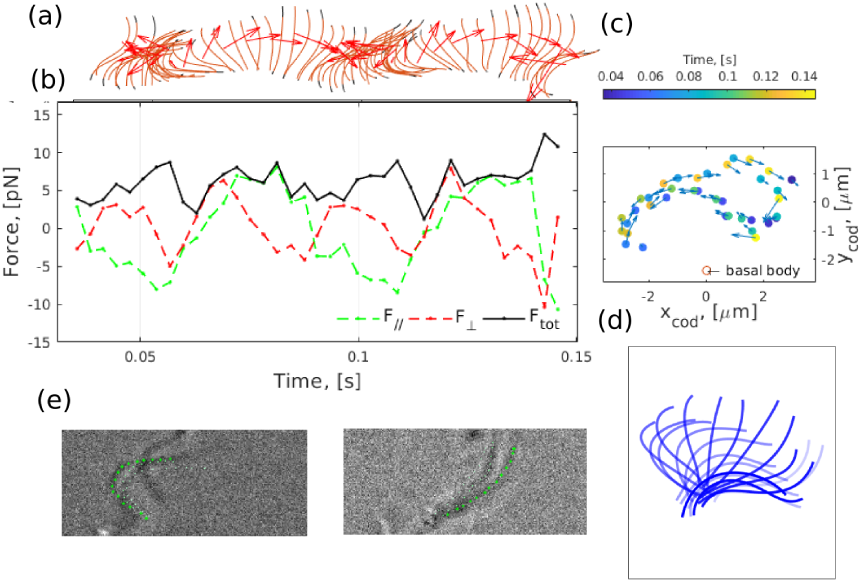
Results from a manual tracking of a isolated cilium. The cilium shape is reconstructed by taking a series of points along the cilium in each image, and then fitting these points with a 4th degree polynomial curve, panel (e). The results of the shapes are in panels (a) and (d). The force exerted by the cilium is calculated using Resistive Force Theory (RFT). Each cilium shape is divided into a series of cylinders 1 *µ*m long. (b) The force that each cylinder is exerting on the surrounding is calculated from the velocity at which it moves, and depends on the orientation of the cylinder (38). The graph shows the force varying with the cycle of beating. (c) The calculated centre of drag of the cilium while beating. The center of drag is calculated as a weighted average of the force along the cilium. The amplitude of the center of drag has been then used as a parameter for the simulations in the main text. Video used for the tracking: http://people.bss.phy.cam.ac.uk/~np451/out/paper/video_ciliumtracking.avi

**Fig. S8.**
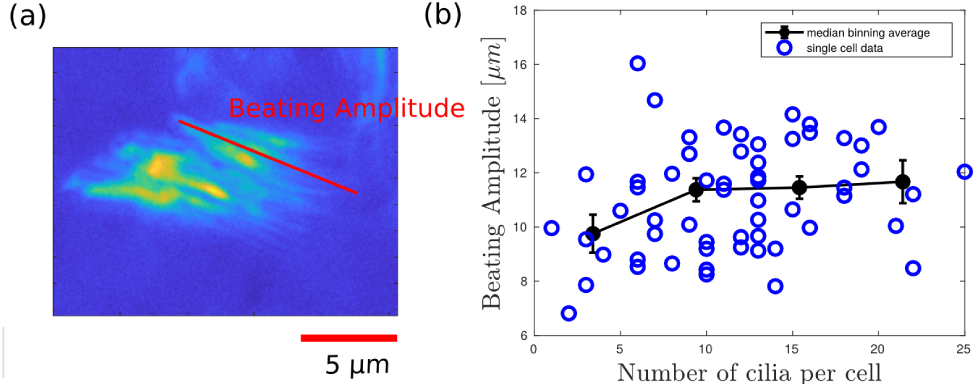
We measured the maximal ciliary beating amplitude for cells with different number of cilia to see if cilia dynamics was affected in these cells. (a) The maximal beating amplitude for a cilium within each cell was measured by inspecting the recordings from top view and marking two extreme points at the power and recovery stroke. For an example, see: http://people.bss.phy.cam.ac.uk/~np451/out/paper/video_amplitudecilia.avi. The length of the red line is the measured amplitude, below there is the standard deviation map of the same cell. It is worth noting that this amplitude is different from the amplitude of the center of drag used for the simulation. (b) The graph shows the beating amplitudes for the 58 cells analysed as a function of the number of cilia per cell and the binning average. Cells with fewer cilia have around 1.5 *µ*m smaller maximal ciliary beating amplitude.

## Notes

There are no conflicts of interest.

## References

1. JR Blake, MA Sleigh, Mechanics of ciliary locomotion. Biol. Rev. 49, 85–125 (1974).

2. E Knight-Jones, Relations between metachronism and the direction of ciliary beat in metazoa. J. Cell Sci. 3, 503–521 (1954).

3. JV Fahy, BF Dickey, Airway mucus function and dysfunction. New Engl. J. Med. 363, 2233–2247 (2010).

4. GR Ramirez-San Juan, et al., Multi-scale spatial heterogeneity enhances particle clearance in airway ciliary arrays. bioRxiv (2019).

5. K Sawamoto, et al., New neurons follow the flow of cerebrospinal fluid in the adult brain. Science 311, 629–632 (2006).

6. M Fliegauf, T Benzing, H Omran, When cilia go bad: cilia defects and ciliopathies. Nat. Rev. 8, 880–893 (2007).

7. B Mitchell, R Jacobs, J Li, S Chien, C Kintner, A positive feedback mechanism governs the polarity and motion of motile cilia. Nature 447, 97 (2007).

8. B Guirao, et al., Coupling between hydrodynamic forces and planar cell polarity orients mammalian motile cilia. Nat. Cell Biol. 12, 341–350 (2010).

9. M Sanderson, M Sleigh, Ciliary activity of cultured rabbit tracheal epithelium: beat pattern and metachrony. J. Cell Sci. 47, 331–347 (1981).

10. S Gueron, K Levit-Gurevich, Energetic considerations of ciliary beating and the advantage of metachronal coordination. Proc. Natl. Acad. Sci. 96, 12240–12245 (1999).

11. HM Oliveira, LV Melo, Huygens synchronization of two clocks. Sci. rep. 5, 11548 (2015).

12. GI Taylor, Analysis of the swimming of microscopic organisms. Proc. Royal Soc. London. 209, 447–461 (1951).

13. A Vilfan, F Julicher, Hydrodynamic flow patterns and synchronization of beating cilia. Phys. Rev. Lett. 96 (2006).

14. J Elgeti, G Gompper, Emergence of metachronal waves in cilia array. Proc. Natl. Acad. Sci. 110, 4470–4475 (2013).

15. B Guirao, J Joanny, Spontaneous creation of microscopic flow and metachronal waves in an array of cilia. Biophys. j. 92, 1900–1917 (2007).

16. T Niedermayer, B Eckhardt, P Lenz, Synchronization, phase locking, and metachronal wave formation in ciliary chains. Chaos 18, 037128 (2008).

17. J Kotar, M Leoni, B Bassetti, M Cosentino Lagomarsino, P Cicuta, Hydrodynamic synchro-nization of colloidal oscillators. Proc. Natl. Acad. Sci. 107, 7669–7673 (2010).

18. N Bruot, P Cicuta, Realizing the physics of motile cilia synchronization with driven colloids. Ann. Rev. Condens. Matter Phys. 7, 1–15 (2016).

19. SL Tamm, TM Sonneborn, RV Dippell, The role of cortical orientation in the control of direction of ciliary beat in paramecium. J. cell biol. 64, 98–112 (1975).

20. H Machemer, Ciliary activity and the origin of metachrony in paramecium: effects of increased viscosity. J. Exp. Biol. 57, 239–259 (1972).

21. M Polin, I Tuval, K Drescher, JP Gollub, RE Goldstein, Chlamydomonas swims with two “gears” in a eukaryotic version of run-and-tumble locomotion. Science 325, 487–490 (2009).

22. DR Brumley, KY Wan, M Polin, RE Goldstein, Flagellar synchronization through direct hydrodynamic interactions. eLife 3, e02750 (2014).

23. DR Brumley, M Polin, TJ Pedley, RE Goldstein, Hydrodynamic synchronization and metachronal waves on the surface of the colonial alga volvox carteri. Phys. Rev. Lett. 109 (2012).

24. G Quaranta, ME Aubin-Tam, D Tam, Hydrodynamics versus intracellular coupling in the synchronization of eukaryotic flagella. Phys. Rev. Lett. 115, 238101 (2015).

25. KY Wan, RE Goldstein, Coordinated beating of algal flagella is mediated by basal coupling. Proc. Natl. Acad. Sci. 113, 2784–93 (2016).

26. KC Leptos, et al., Antiphase synchronization in a flagellar-dominance mutant of chlamydomonas. Phys. Rev. Lett. 111, 158101 (2013).

27. Y Liu, R Claydon, M Polin, DR Brumley, Transitions in synchronization states of model cilia through basal-connection coupling. J. R. Soc. Interface 15, 20180450 (2018).

28. GS Klindt, C Ruloff, C Wagner, BM Friedrich, In-phase and anti-phase flagellar synchronization by waveform compliance and basal coupling. New J. Phys. 19, 113052 (2017).

29. A Mahuzier, et al., Ependymal cilia beating induces an actin network to protect centrioles against shear stress. Nat. Comms. 117, 094101 (2017).

30. EK Vladar, RD Bayly, AM Sangoram, MP Scott, JD Axelrod, Microtubules enable the planar cell polarity of airway cilia. Curr. Biol. 22, 2203–2212 (2012).

31. ME Werner, et al., Actin and microtubules drive differential aspects of planar cell polarity in multiciliated cells. J. Cell Biol. 195, 19–26 (2011).

32. N Delgehyr, et al., Ependymal cell differentiation, from monociliated to multiciliated cells in Meth. Cell Biol. (Elsevier) Vol. 127, pp. 19–35 (2015).

33. R Faubel, C Westendorf, E Bodenschatz, G Eichele, Cilia-based flow network in the brain ventricles. Science 353, 176–178 (2016).

34. N Bruot, L Damet, J Kotar, P Cicuta, M Cosentino Lagomarsino, Noise and synchronization of a single active colloid. Phys. Rev. Lett. 107, 094101 (2011).

35. A Pikovsky, M Rosenblum, J Kurths, Synchronization: a universal concept in nonlinear sciences. (Cambridge university press) Vol. 12, (2003).

36. S Dimova, et al., High-speed digital imaging method for ciliary beat frequency measurement. J. pharm. pharmacol. 57, 521–526 (2005).

37. JH Sisson, J Stoner, B Ammons, T Wyatt, All-digital image capture and whole-field analysis of ciliary beat frequency. J. microscopy 211, 103–111 (2003).

38. E Lauga, TR Powers, The hydrodynamics of swimming microorganisms. Rep. Prog. Phys. 72, 096601 (2009).

39. C Allain, M Cloitre, The effects of gravity on the aggregation and the gelation of colloids. Adv. colloid interface science 46, 129–138 (1993).

40. AJ Hunt, F Gittes, J Howard, The force exerted by a single kinesin molecule against a viscous load. Biophys. journal 67, 766–781 (1994).

41. GS Klindt, C Ruloff, C Wagner, BM Friedrich, Load response of the flagellar beat. Phys. review letters 117, 258101 (2016).

42. N De Mestre, W Russel, Low-reynolds-number translation of a slender cylinder near a plane wall. J. Eng. Math. 9, 81–91 (1975).

43. K Drescher, N Goldstein, Raymond E and Michel, M Polin, I Tuval, Direct measurement of the flow field around swimming microorganisms. Phys. Rev. Lett. 105, 168101 (2010).

44. E Hamilton, N Bruot, P Cicuta, The chimera state in colloidal phase oscillators with hydrodynamic interaction. Chaos 27, 123108 (2017).

45. E Hamilton, P Cicuta, Interpreting the synchronisation of driven colloidal oscillators via the mean pair interaction. New J. Phys. 20, 093028 (2018).

46. M Cosentino Lagomarsino, P Jona, B Bassetti, Metachronal waves for deterministic switching two-state oscillators with hydrodynamic interaction. Phys. Rev. E 68, 021908 (2003).

47. W C. H Stark, Metachronal waves in a chain of rowers with hydrodynamic interactions. Eur. Phys. J. E 34 (2011).

48. N Bruot, P Cicuta, Emergence of polar order and cooperativity in hydrodynamically coupled model cilia. J. R. Soc.: Interface 10, 20130571 (2013).

49. B Button, et al., A periciliary brush promotes the lung health by separating the mucus layer from airway epithelia. Science 337, 937–941 (2012).

50. D Trieu, TK Waddell, AP McGuigan, A microfluidic device to apply shear stresses to polarizing ciliated airway epithelium using air flow. Biomicrofluidics 8, 064104 (2014).

51. B Chueh, et al., Leakage-free bonding of pourous membranes into layered microfluidic array system. Anal. Chem. 79, 3504–3508 (2007).

52. KH Benam, et al., Small airway-on-a-chip enables analysis of human lung inflammation and drug responses in vitro. Nat. methods 13, 151 (2015).

53. A Tourovskaia, X Figueroa-Masot, A Folch, Differentiation-on-a-chip: a microfluidic platform for long-term cell culture studies. Lab Chip 5, 14–19 (2005).

54. L Kim, YC Toh, J Voldman, H Yu, A practical guide to microfluidic perfusion culture of adherent mammalian cells. Lab Chip 7, 681–694 (2007).

55. M Ryser, A Burn, T Wessel, M Frenz, J Rička, Functional imaging of mucociliary phenomena. Eur. Biophys. J. 37, 35–54 (2007).

